# Many ways to stick the landing: novel righting strategies allow spotted lanternfly nymphs to land on diverse substrates

**DOI:** 10.1101/2021.04.12.439561

**Authors:** Suzanne Amador Kane, Theodore Bien, Luis Contreras-Orendain, Michael F. Ochs, S. Tonia Hsieh

## Abstract

Unlike large animals, insects and other very small animals are so unsusceptible to impact-related injuries that they can use falling for dispersal and predator evasion. Reorienting to land upright can mitigate lost access to resources and predation risk. Such behaviors are critical for the spotted lanternfly (SLF) (*Lycorma delicatula*), an invasive, destructive insect pest spreading rapidly in the US. High-speed video of SLF nymphs released under different conditions showed that these insects self-right using both active midair righting motions previously reported for other insects, and novel post-impact mechanisms that take advantage of their ability to experience near-total energy loss on impact. Unlike during terrestrial self-righting, in which an animal initially at rest on its back uses appendage motions to flip over, SLF nymphs impacted the surface at varying angles and then self-righted during the rebound using coordinated body rotations, foot-substrate adhesion, and active leg motions. These previously-unreported strategies were found to promote disproportionately upright, secure landings on both hard, flat surfaces and tilted, compliant host plant leaves. Our results highlight the importance of examining biomechanical phenomena in ecologically-relevant contexts, and show that, for small animals, the post-impact bounce period can be critical for achieving an upright landing.

## 1. Introduction

Falling is a frequent and unavoidable fact of life for animals in a wide range of environments. In response, many climbing arthropods and arboreal vertebrates have evolved a variety of strategies to help them land safely, such as gliding (1), parachuting, and righting (i.e., reorienting so as to land upright) (2). Although smaller organisms are not at direct risk from impact-related injury (3), landing upright can still maximize survival by minimizing the metabolic cost of terrestrial righting, facilitating predator evasion (4), and mitigating other risks (e.g., hunger, desiccation, habitat and territory loss, etc.) (5). Because dropping is also a strategy used by animals for dispersal (6) and predator avoidance (4), understanding these behaviors has a wide variety of implications for ecology, as well as providing inspiration for robotics (7).

Among insects and other arthropods, righting behaviors have been categorized into two broad groups: aerial righting and terrestrial righting (8). Aerial righting consists of body reorientation during the fall, and typically includes an active push off of the surface with the limbs imparting an initial rotation on the body. Some small arthropods use a stereotypical falling body posture to take advantage of aerodynamic drag on the body and legs for aerial righting and even maneuvering during gliding, as found for pea aphids, stick insect instars, canopy ants and spiders (2,9–11). Repositioning of various body parts can also facilitate reorientation to an upright posture and a controlled landing, ideally with feet in contact with the substrate. Just as some larger, flexible vertebrates (e.g., cats, rabbits, squirrels, lizards) tend to use a combination of body, limb, and tail inertia to right themselves while falling (2), similar strategies appear to be used among falling stick insect nymphs, which have a relatively flexible and long body (12). On the other hand, terrestrial righting consists of determining how an animal that is on its back can get back onto its feet. Among insects, this usually involves a period of pushing off of the substrate using a combination of legs, wings, and the body imparting a rocking motion on the body until a leg can gain enough purchase to complete an upright flip, as observed in locusts and cockroaches (7,13,14).

We elected to study falling and righting in the spotted lanternfly (*Lycorma delicatula*) (SLF), a phloem-feeding planthopper native to China and south Asia that has become a major invasive pest threatening agriculture and forestry in the US since its introduction in 2014 (15), making research that can inform the development of more effective traps and other deterrents particularly urgent (16). The spotted lanternfly undergoes a rapid lifecycle, quadrupling in length (15) and progressing through four wingless nymphal stages (instars) in three to four months before emerging as a winged adult (17). (Fig. 1A,B) Although nymphs and adults are able to cling securely to leaflets, stems, branches and other surfaces using a combination of tarsal claws and adhesive pads (arolia) (18,19), they frequently drop out of trees and climb back into the canopy of the same or nearby trees in response to obstacles, wind, or predator attack (20). In light of their rapid growth, the metabolic cost of frequent climbing and interruptions in time spent feeding likely impose significant fitness costs. This raises the question of whether falling SLF nymphs are able to land securely on lower leaves of their host plant. The SLF’s preferred host, the *Ailanthus altissima* tree, has dense layered foliage consisting of pinnately compound leaflets (Fig. 1C, D) that likely offer numerous landing targets for falling insects. Our observations indicate that SLFs that either drop or jump often land on underlying or neighboring plants. Consistent with this, capture-mark-recapture studies have shown that SLFs frequently remain on or nearby a healthy host *A. altissima* tree (21). We therefore hypothesized that this species should exhibit righting during dropping onto leaves and the ground. As far as we are aware, biomechanical studies of SLF nymphs have measured their walking, jumping and climbing ranges and rates, not their behavior during dropping or their righting capabilities (6,22).

**Figure 1.**
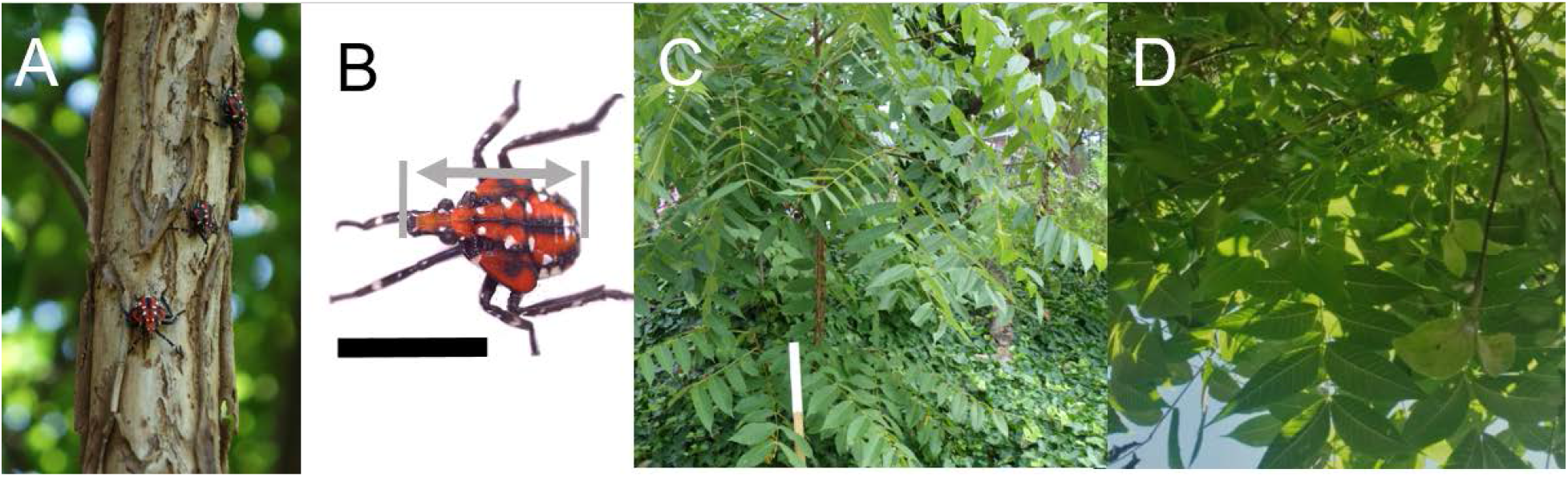
A) Fourth instar spotted lanternfly nymphs on a trunk. B) Close-up of a fourth instar spotted lanternfly nymph showing our definition of body length (gray arrow) (black line = 10 mm scale bar). Photographs of the spotted lanternfly’s preferred native host tree, *A. altissima*, showing (C) the release distance (white bar = 200 mm) used in most experiments in this study and (D) a view from the ground looking upward into the canopy, showing how the densely overlapping leaflets offer many landing opportunities for falling nymphs.

In this study, we addressed the following research questions via a series of laboratory experiments on SLF fourth instar nymphs. First, we sought to quantify whether spotted lanternfly nymphs indeed do self-right more often than expected by chance when dropped. Once confirmed, we examined the strategies they used for righting and landing in general. Finally, we asked whether these righting behaviors influence their ability to land on lower layers of foliage, to avoid completely falling out of the tree.

## 2. Methods

Live, fourth instar spotted lanternfly nymphs were collected and studied within a quarantine zone in southeastern Pennsylvania, US (40.006525, −75.256714) in July-August 2020. All experiments were performed indoors in still air at 24 ± 3 deg C. Nymphs were collected by hand or using an insect net and scoop-shaped forceps from natural habitats, primarily *A. altissima* trees. Only intact, healthy, and active insects were studied. Fourth instar nymphs were identified by their distinctive red, black and white coloration. (Figs. 1A, 2A) Specimens not immediately used in experiments were stored in a sealed container with freshly picked *A. altissima* foliage and wet tissues. Insects maintained in this way retained their normal levels of activity for at least 48 hours. Because this species is the subject of an eradication program (23), all specimens were euthanized by freezing after experimentation. For studies of dead specimens, we used frozen insects that were thawed and either used within 30 min of thawing or stored in 49% relative humidity chambers to avoid desiccation and to preserve their native biomechanical properties (24). Specimen body length, *L* = 11.8 mm [10.3, 12.6] mm (mean, range) (Fig. 1B) was measured to ± 0.05 mm either using digital calipers (model SV-03-150, E-base Measuring Tools, Yunlin, Taiwan) or using the *measure* function in ImageJ (25) on digital photographs including a mm-ruled scale. Body masses, *m* = 66 ± 18 mg, [40, 100] mg (N = 16, mean ± SD, range), were measured to ± 0.4 mg with an analytical balance (Explorer, Ohaus, Parsippany, NJ US). (See S1 Table for morphometric data).

### 2.1 Video studies of dropping and landing experiments

We performed a variety of experiments on SLF nymphs filmed during dropping to determine their midair body motions and center of mass trajectories while falling and their landing behavior. Most high-speed videos of spotted lanternfly specimens were filmed using an SA-3 high-speed camera (Photron, Tokyo, Japan) (monochrome, 1024 × 1024 pixel resolution, 1000 frames/s; exposure 500 microsec) illuminated by a Nila Zaila LED light (Nila, Inc. Altadena, CA). For filming SLFs releasing voluntarily from surfaces and falling on leaves, we used a color Chronos 1.4 camera at a higher frame rate (800 × 600 pixels, 2837 fps; exposure 343 microsec, Krontech, Burnaby, BC, Canada) and LED light source (SL-200W, Godox, Shenzhen, China). A second perspective was provided by a mirror included in the field of view to allow visualization of body pose and rotational behavior.

Dropping experiments were performed on live specimens and on dead SLF nymphs with their legs contracted close to the body (“dead/tucked”). An additional set of dead nymphs (“dead/spread”) were pinned with their legs spread and fixed with a small drop of cyanoacrylate glue applied using a fine needle to each leg joint (total added mass 1.0 ± 0.2 mg; 1-2% body mass), so as to position the forelegs above the dorsal plane to approximate the posture we observed for SLFs, which was similar to those reported for falling pea aphids (11). Glue was applied so as to avoid coating the feet, which still adhered to substrates after being glued. Following previous studies (5,11,26), we released specimens with the goal of achieving low initial speed and spin while controlling initial falling orientation. Live and dead specimens were released using featherweight entomology tweezers (DRENTF01, DR Instruments, Palos Hills, IL US) (Fig. 2A) oriented side, head down or caudal end down, similar to methods used in (11,27). To release specimens upright or upside-down, we grasped them initially with fingers on the sides of the scutellum. If a nymph responded to handling by feigning death, it was breathed on until it spread its legs and moved actively. All specimens were inspected after manipulation during experiments and showed no effect of handling. Live specimens sometimes were measured in a second session after being marked in one or two places with white paint (0.9 ± 0.2 mg; 1-2% body mass); this added mass is unlikely to influence specimen motility given that heavier (6.5% body mass) harmonic radar tags do not significantly affect SLF fourth instar nymph walking, climbing, jumping or survivorship (22). While we tried to achieve a balanced study design, the tendency of this species to fatigue rapidly limited our ability to conduct a uniform number of trials on each specimen.

**Figure 2.**
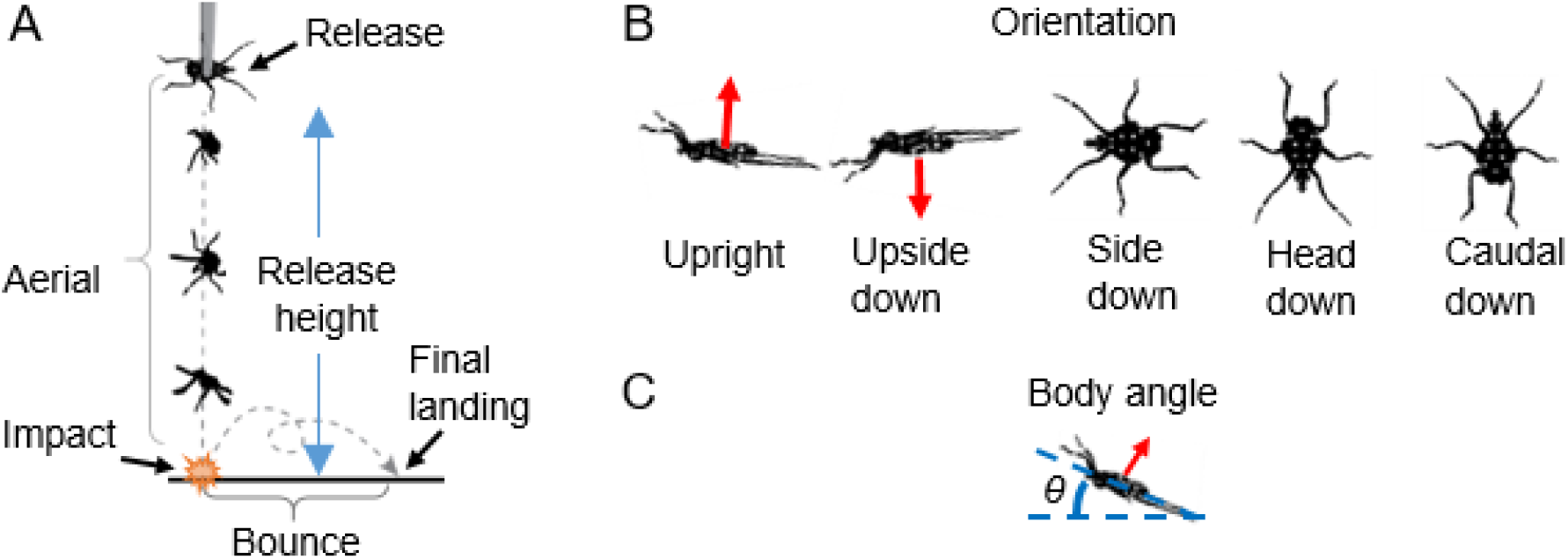
A) Schematic of the dropping experiment, showing how the five different phases of motion used to analyze the outcomes were defined. B) Illustration of the five orientations used to describe releases, impacts and final landings. (Red arrows indicate dorsoventral axis.) C) Geometry used to define the cranial-caudal body axis angle, *θ*, relative to horizontal.

Specimens were dropped artificially (i.e., either from tweezers or manually) from a uniform height of 200 mm, approximately 17 body lengths, measured using a nearby vertical ruler. Preliminary tests established that this range was high enough for a large fraction of specimens to land upright. This height allowed comparison with previous research on righting by pea aphids that used a similar range of heights (11) and by stick insect nymphs (12), which self-righted aerially over a similar height in terms of body lengths. We also note that this choice of falling distance lies in the range of distances between neighboring leaflets in *A. altissima* trees. (Fig. 1C)

For video studies, specimens fell onto one of two landing substrates: 1) a hard, horizontal surface covered with white, art-quality watercolor paper; 2) a freshly-harvested, freely-suspended *A. altissima* leaf taped to a post such that the surfaces of its freely-suspended leaflets were inclined by [0,30] deg relative to horizontal. SLF nymphs were observed to be able to achieve a secure footing by using their tarsal claws and arolia on both hard and leaf substrates. Because it was difficult to achieve a reproducible impact location on the leaflets, the leaf substrate experiments provided insight only into impact and post-impact behaviors that resulted in a successful landing.

To study whether SLF nymphs able to launch voluntarily from surfaces exhibited different behaviors during falling and landing, we also filmed SLF nymphs that were stimulated to release from the wall of a clear acrylic box by moving a plastic insect toward them or gently breathing on them—a trigger we observed to elicit dropping behavior in the field. Because of the known tendency of these insects to climb (6), we were unable to control the drop distance in these trials to agree with that used for the artificially released specimens.

We divided each falling trial into four periods for analysis: aerial, impact, post-impact, and final landing. (Fig. 2A) Specimen orientation at initial impact and final landing (i.e., after coming to rest post-impact) was scored by frame-by-frame video analysis into the best agreement with five categories similar to definitions used in (11). (Fig. 2B) In the following, a body axis is referred to as “horizontal” and “vertical” when it agrees with the respective direction within 45 deg; e.g., a horizontal cranial-caudal axis corresponds to *θ* ≤ ±45 deg. (Fig. 2C) With this convention, specimens oriented “upright” or “upside-down” had their dorsal or ventral side uppermost, respectively, and horizontal cranial-caudal and medio-lateral axes. The “side” orientation had a horizontal cranial-caudal angle and vertical medio-lateral axis with the left side oriented downward, while the “head” and “caudal” orientations had a vertical cranial-caudal axis and the head or caudal side oriented downward, respectively. We also recorded whether the nymphs bounced during landing, defined as vertical motion of the body center of mass after impact in which at least two feet lost contact with the ground.

### 2.2 Image analysis

Videos were analyzed using custom image analysis code written in MATLAB v2020A with the machine vision and curve fitting toolboxes (Mathworks, Natick MA) (Supplemental Materials); all italicized functions referenced below are from MATLAB unless noted otherwise. The MATLAB camera calibrator was used to calibrate and correct each camera for lens distortion before analysis (mean reprojection error: ≤ 0.3 pixel). The spatial calibration was measured from images of a ruler at the same distance as the specimen (range [1.7, 5.8] pixel/mm), and checked using known body dimensions. The maximum bounce height (defined as the difference between the body midpoint at the lowest and highest heights immediately after impact) was measured manually using ImageJ (25). For automated tracking of specimens, all images were blurred using a Gaussian filter using *imgaussfilt* with sigma of 0.6 mm to reduce noise. To isolate the specimen’s image from background, we then computed a background image by taking the median intensity of the video using *median2*, and subtracted this background image from each video frame using *imabsdiff* to compute the absolute difference between the two images. The difference image was contrast-enhanced using *adapthisteq* to correct for nonuniform illumination, and thresholded using *imbinarize* to create a binary image of a white specimen on a black background. If necessary, the morphological command *imclose* was used to fill in holes on the specimen due to white spots. For tracking and determining body orientation, the resulting binary image was processed using the morphological operation *imopen* (a dilation followed by an erosion over approximately 1.5 mm) to remove the legs, after which *regionprops* was used compute the body centroid (*x*, *y*) and an ellipse that has the same normalized second central moments as the body. The orientation angle, *θ*, of the body (i.e., the angle of the cranial-caudal axis relative to horizontal) was tracked using a combination of the orientation of the body ellipse’s major axis and the angle at which the distance between the body’s outline and its centroid is at a maximum due to the protruding head.

### 2.3 Statistical methods and data analysis

The statistical analysis of data was performed with R v3.6.3 (28). Outcomes for experiments were analyzed using Fisher’s exact tests or χ^2^ tests. Where Fisher’s exact test sample size exceeded computing capacity, simulated p-values were generated from 2000 simulations. All results are reported as mean [95% C.I.] unless noted otherwise. ANOVA with Tukey’s Honest Significant Differences (HSD) was used to determine p-values between different conditions for kinematics measures such as speeds and bounce heights. Data and code required to reproduce all results are included in the Supplementary Materials.

The coordinate data from video tracking were analyzed without further processing using nonlinear least squares fitting in MATLAB. The vertical falling trajectory (*y* vs time, *t*), was fit to the equation of motion for the case of a drag force quadratic in speed:

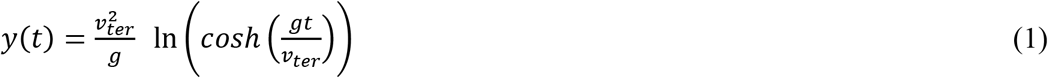

where *v_ter_* is terminal speed and *g* = 9.81 m/s^2^ is the acceleration of gravity. The horizontal data (*x* vs *t*) were fit to a quartic polynomial (the lowest order polynomial found to result in mean fit residuals < 0.8 mm). Goodness-of-fit was assessed using *R^2^* and residuals analysis. The speed before impact, *v_imp_*, and terminal speed, *v_ter_* were determined from fit parameters, and then used to compute Reynolds number, *Re* = *L v /* υ, where *L* = body length and the kinematic viscosity of air, υ = 15.34 × 10^−6^ m^2^/s (29), as well as the fractional collisional energy loss on impact, 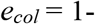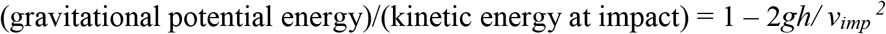, where *h* = maximum bounce height after impact.

We were also interested in measuring whether the rotation rate of the specimen about the normal to the image plane varied during the fall. This quantity is relevant because conservation of angular momentum dictates that the rotation rate about any given axis is constant for constant specimen rotational inertia and zero net torque along that direction (30). Consequently, in order for its rotation rate to vary throughout the fall, the specimen must either experience nonzero torque due to aerodynamic drag or move its legs so as to vary rotational inertia. Using MATLAB, we manually measured the specimen’s average aerial rotation rate, *Ω* = Δ*θ /*Δ*t*, from the change in the body angle on the image, Δ*θ*, between two frames recorded Δ*t* = 25 ms apart. The associated measurement uncertainty was determined from the error in determining the initial and final orientations of the specimen’s cranial-caudal axis on video. The initial rotation rate, *Ω_rel_*, was measured shortly (50 ms) after release, to ensure that the specimen was clear of the tweezers or wall. To determine if *Ω* varied throughout the fall, the rotation rate also was measured at the approximate midpoint of the fall (125 ms after release), *Ω_mid_*, and immediately before impact, *Ω_imp_*.

## 3. Results

### 3.1 Effect of orientation at release on landing

We analyzed high-speed video of falling and landing on hard surfaces for five trials for each of five release orientations from tweezers for five different live SLF nymphs (125 trials total). To determine whether release orientation impacted the distribution of orientations on impact and final landing, we considered three orientation outcomes for impact: upright, upside-down and other (comprising side, caudal and head down) and two for landing: upright and upside-down. (Fig. 3) We found that neither the orientation distribution on impact nor on final landing showed significant differences based on release orientation (orientation at impact: Fisher’s exact test, p = 0.22; orientation at final landing, χ^2^ test, p = 0.80). This suggests that orientation upon impact and landing are independent of release orientation. Similarly, for dead nymphs (30 specimens, 1 trial each per release orientation) dropped onto a hard substrate, the distributions of final landing orientations did not depend significantly on release orientation for spread legs (χ^2^ test, p = 0.86). (S1 Dataset) We consequently analyzed these datasets summed over release orientations, and only recorded data for a single release orientation (side down) when studying dead specimens with legs spread and tucked.

**Figure 3.**
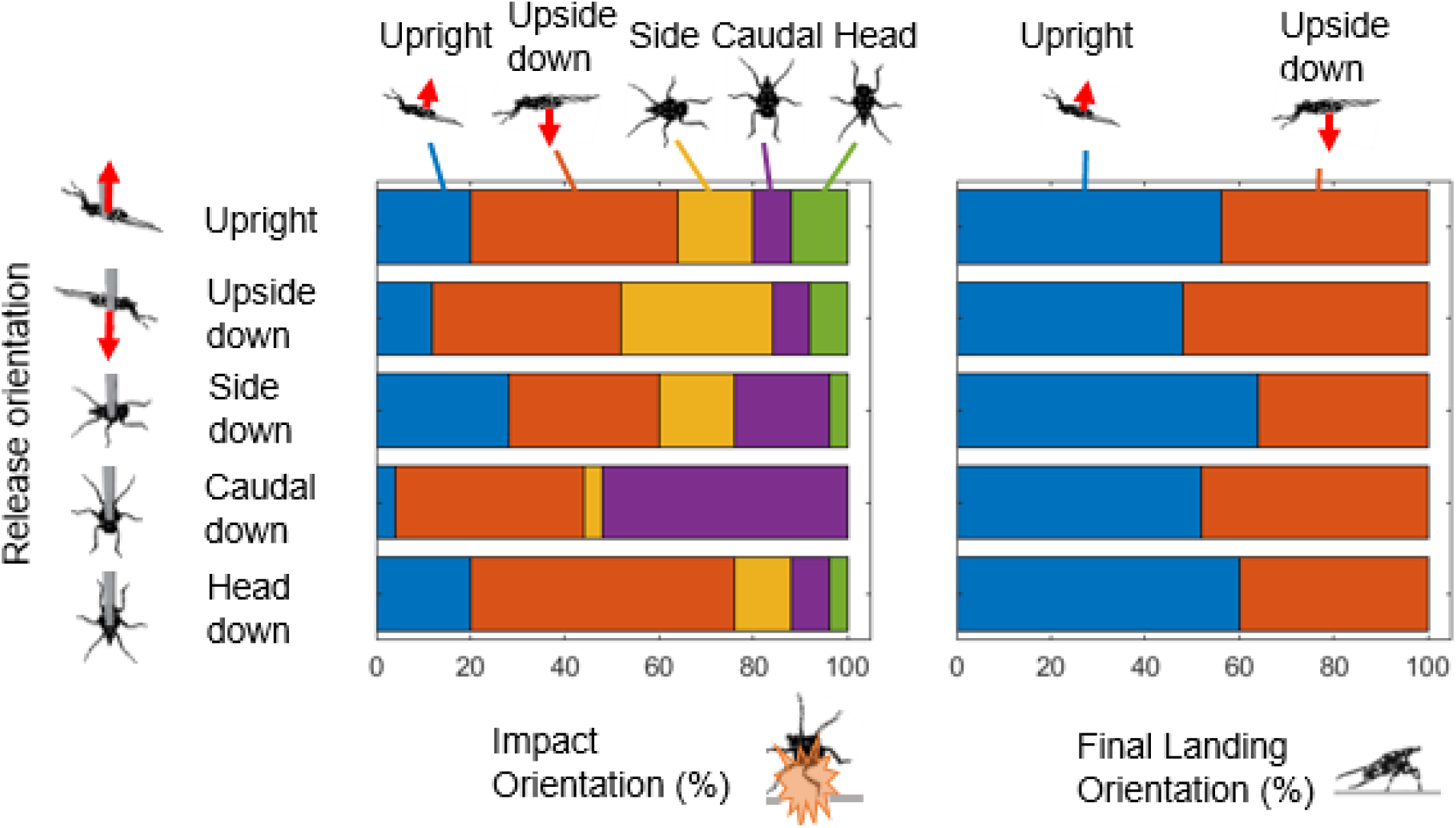
Distribution of orientation at impact and final landing for live spotted lanternfly fourth instar nymphs dropped on a hard surface from each of five release orientations. (Red arrow points towards the dorsal surface when viewed in lateral aspect.)

### 3.2 Aerial phase

Fig. 4A shows a typical sequence of motions by live SLF nymphs during the aerial phase of the dropping experiments. ([S1Movie](https://www.youtube.com/watch?app=desktop&v=Wd-3L8Az7SQ&feature=youtu.be))In the majority of trials (97.1% = 135/139) artificially released specimens assumed a stereotypical falling posture within 0.079 s [0.029, 0.129] s after release, in which they spread their legs fully and held them slightly above the dorsal plane until impact. (Fig. 4B) Results from kinematic data analysis are shown as summary statistics in Table 1. All measured trajectories of artificially released live and dead SLF nymphs were predominantly vertical (mean horizontal excursions ≤ 4.3 mm). We were successful at filming a total of 15 voluntary release trials for five specimens from 354 mm [342, 365] mm above the hard substrate (Fig. 5B). These trajectories displayed greater horizontal excursions (24 mm [13, 35] mm) than observed for the artificial releases. All falling trajectories for all conditions agreed well with a quadratic drag model (R^2^ ≥ 0.9998; fit-residuals 0.9 mm [0.7, 1.0] mm). (Fig. 4C) Terminal speed did not differ significantly between live specimens released voluntarily and artificially (Tukey’s HSD, p = 0.48) (Fig. 4D). Dead specimens with their legs tucked had a mean terminal velocity that was greater and statistically different from all other conditions (i.e., live and dead with legs spread) (Tukey’s HSD, p < 0.0001), whereas analysis of live and dead with legs spread found no significant differences (Tukey’s HSD, p > 0.74). (Fig. 4D). During falling, all SLF nymphs studied here had speeds corresponding to Reynolds number *Re* in the range, [10^2^, 10^4^], consistent with values reported for gliding arthropods (29).

**Table 1.**
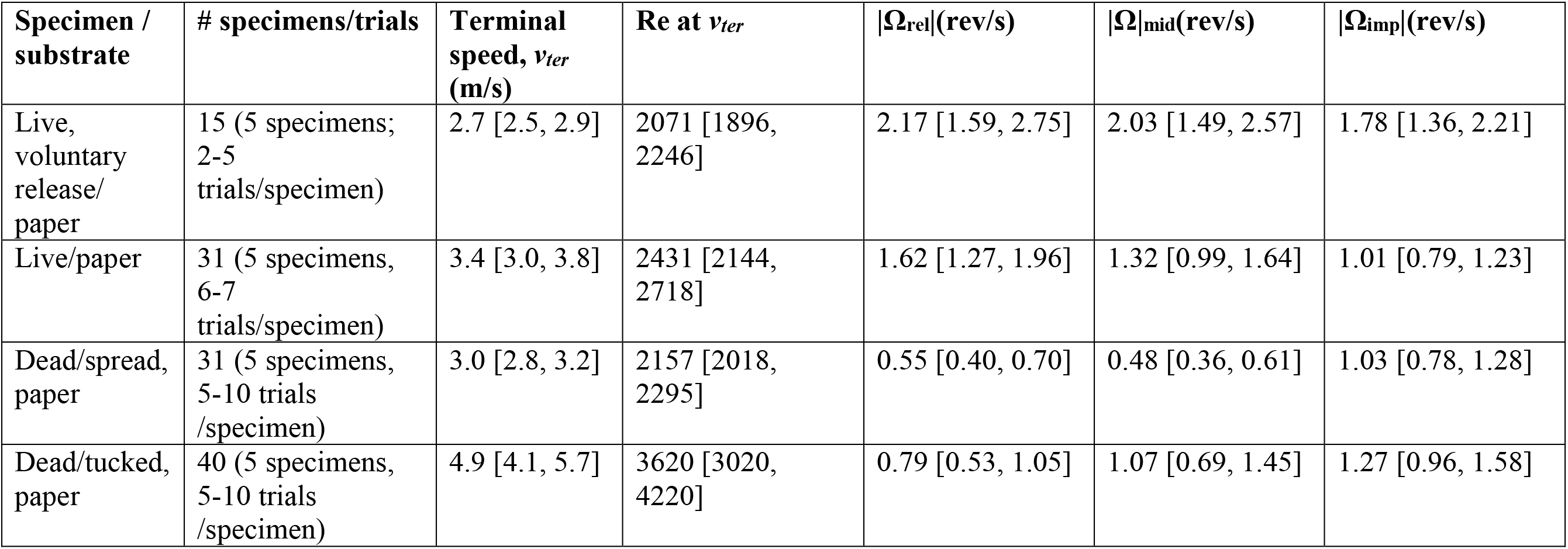
Summary statistics for spotted lanternfly fourth instar (N4) nymph aerial kinematics.

**Figure 4.**
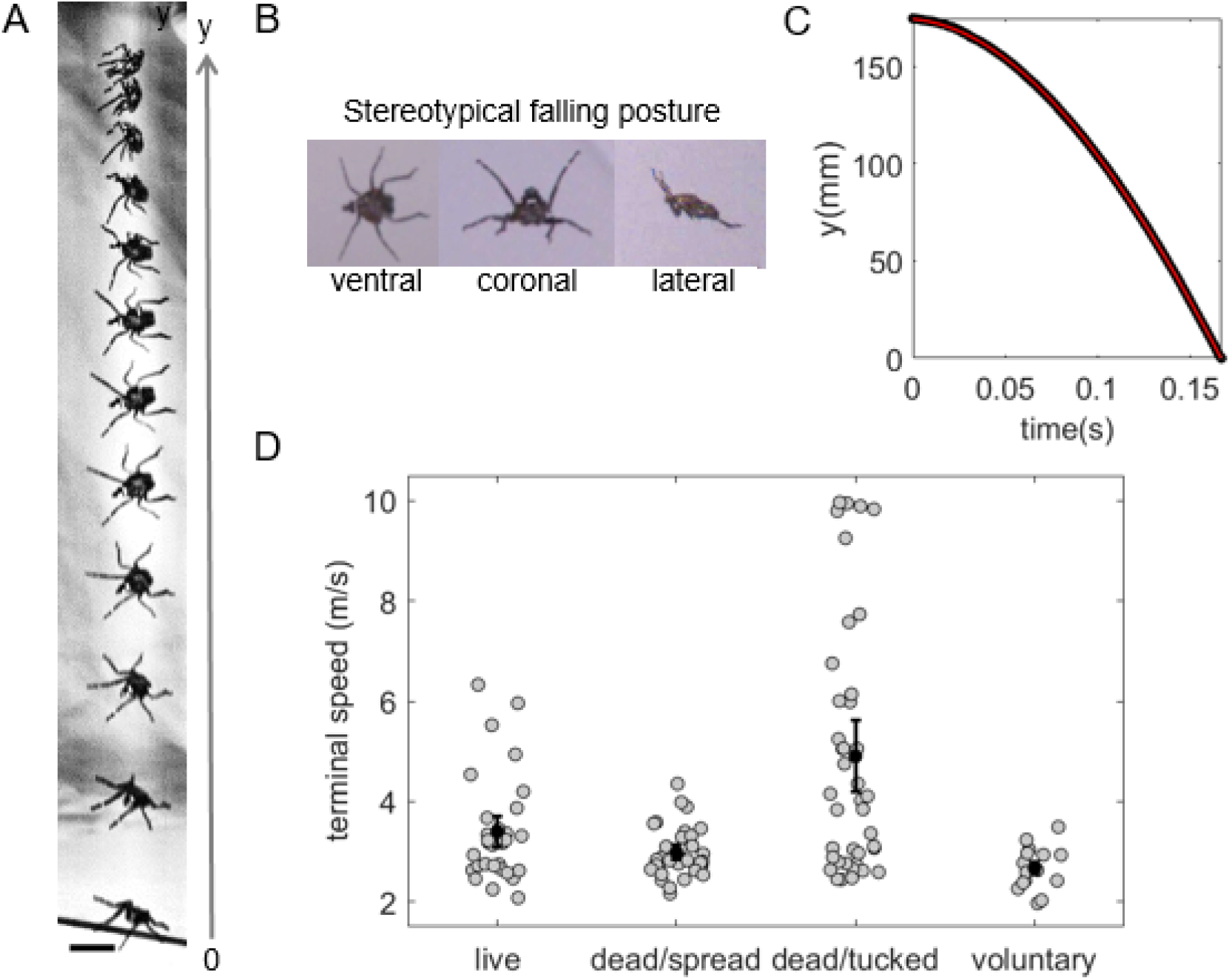
A) Stereotypical falling posture assumed by spotted lanternfly nymphs after dropping. B) Superimposed sequence of video frames recorded every 15 ms showing a spotted lanternfly nymph falling 200 mm. (Scale bar: 10 mm). C) Measured (open circles) and fitted (red line) vertical position, *y*, of the specimen shown in B) plotted vs time. D) Fitted terminal falling speed distributions for live and dead specimens artificially dropped from 20 cm and live specimens voluntarily releasing from 35 cm. (black circles: mean; error bars: 95% CI; gray circles: all data, jittered for visibility.)

**Figure 5.**
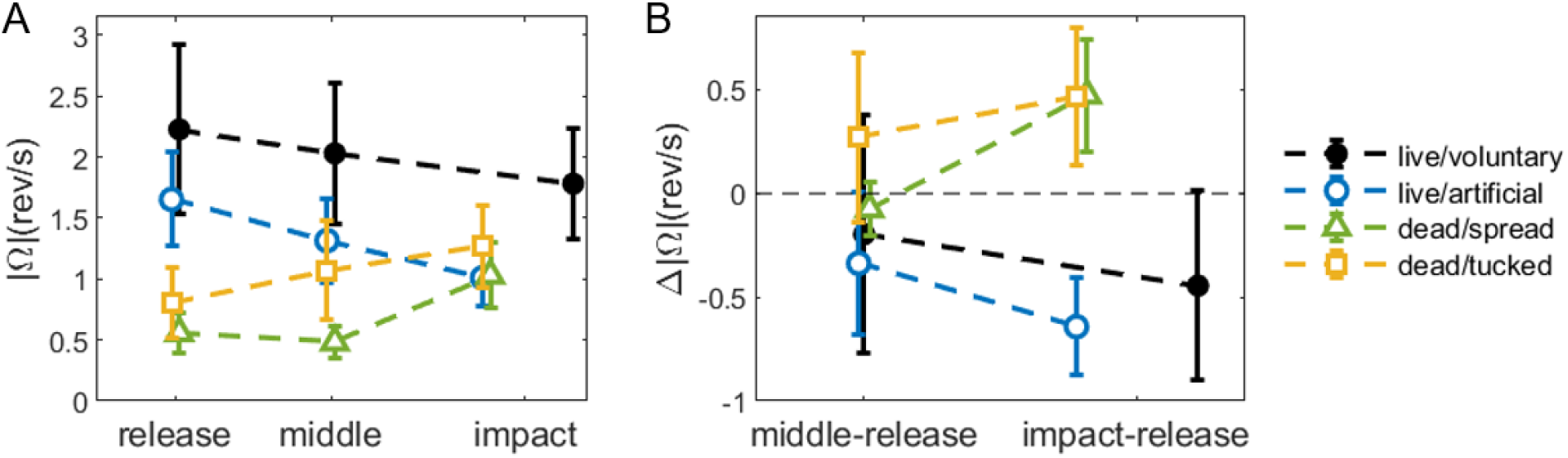
Rate of spotted lanternfly nymph body rotation in the image plane upon release, at an approximate midpoint during the fall, and immediately before impact. (A) Rotation rate magnitudes show different trends during the fall period for live nymphs (circles) relate to dead specimens with legs spread (triangles) or tucked (squares). (B) Plots of the change in rotation rate magnitude between the midpoint and at impact relative to release (equivalent to scaling the initial rotation rate at release to zero) for different release methods and specimen preparations.). (Error bars show 95% CI, which were similar to instrumental measurement uncertainties. Horizontal distances between data points are proportional to time; data also are jittered for visibility.)

We next consider results for the aerial rotational kinematics, which differed significantly between live and dead specimens. The rotation rate after release, *Ω_rel_*, of live specimens was significantly larger than for dead specimens (one-sided t-test p = 2.4 × 10^−5^ vs dead with legs spread and p = 0.0013 vs dead with legs tucked). (Fig 5A) The value of *Ω_rel_* was independent of the release method for live specimens, and of pose for the dead specimens (two-sided t-test, p = 0.365 live by release method, p = 0.157 dead by pose). To illustrate how rotation rate changed during falling, we computed the change in rotation rate magnitude, Δ|*Ω|*, at the midpoint and impact relative to its value at release (Fig. 5B). These data showed that rotation rates tend to decrease with fall time among live specimens, but increase or stay the same for the dead specimens. In some cases, live nymphs changed the direction of their rotation or increased their spin to a greater rotation rate mid-fall than at either release or impact. A variety of related behaviors could be observed on some videos of live nymphs: 1) pushing off the wall (voluntary) and tweezers (artificial) so as to impart an initial spin; 2) changing the orientation and extension of their legs during falls so as to alter their rotational inertia ([S1Movie](https://www.youtube.com/watch?app=desktop&v=Wd-3L8Az7SQ&feature=youtu.be)).

### 3.3 Impact orientation

Fig. 6 shows the orientation distributions for specimens at impact and final landing on the hard substrate for live and dead SLF nymphs for artificial and voluntary releases. The impact orientation was found to differ between live nymphs and dead with legs tucked (Fisher exact test, p = 0.00005) but not between live and dead with legs spread (Fisher exact test, p = 0.95). For voluntary releases, a greater fraction impacted upright (67%) than when artificially released, and no specimens that released voluntarily impacted upside down (Fig 6B). The distributions of orientations on impact differed significantly for live specimens between the two release conditions (Fisher’s exact test, p < 8 × 10^−6^ and p = 0.00039, impact and 200 mm respectively). (Fig. 6) To make sure that release height did not influence this last finding, we also measured the distribution of orientations after specimens that released voluntarily had fallen 200 mm, the height used for artificial releases. This distribution also was similar to that found at impact (Fisher’s exact test, p = 0.71). (Fig. 6B)

**Figure 6.**
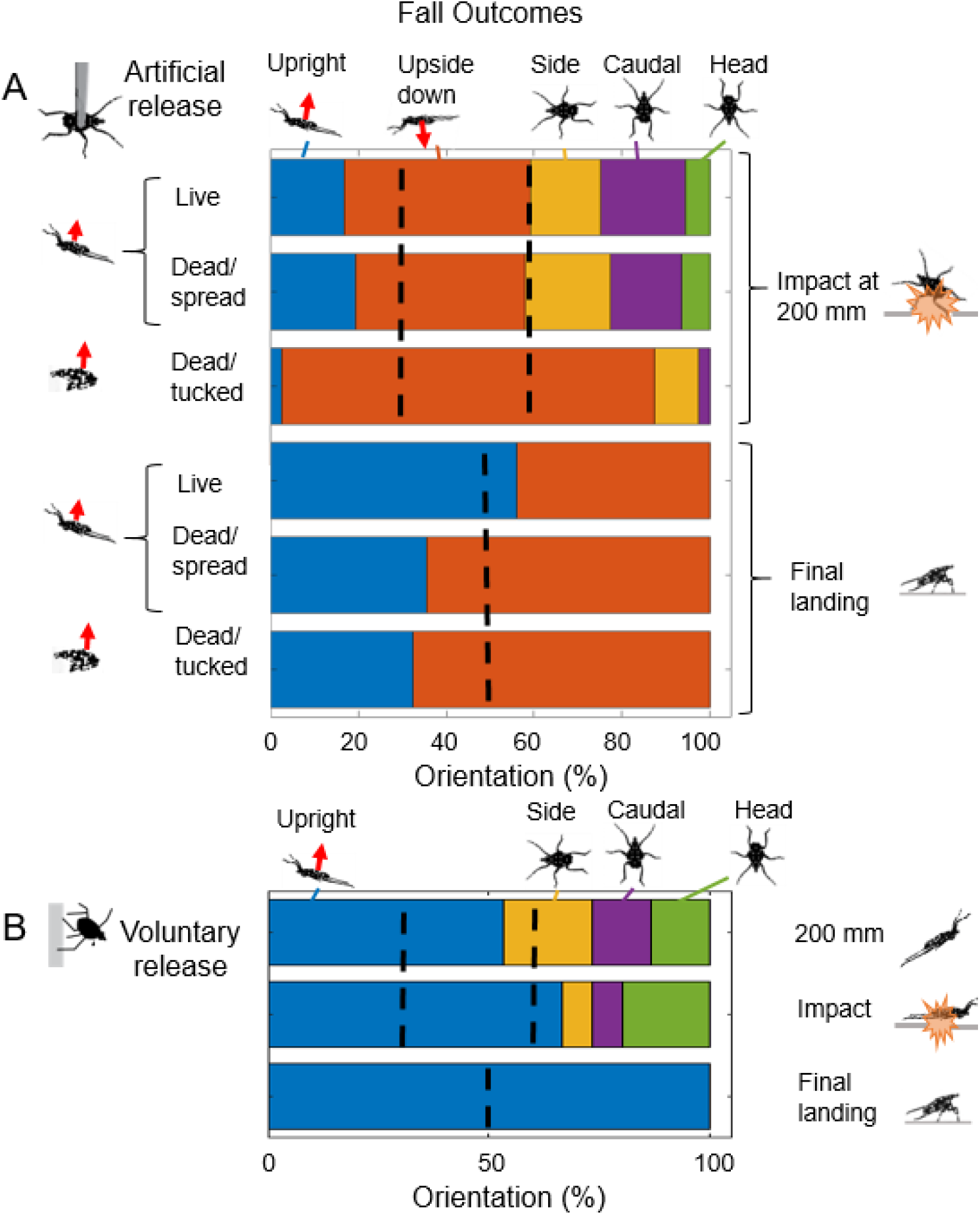
Distribution of impact and final landing orientations for spotted lanternfly nymphs (A) dropped artificially from tweezers and (B) releasing voluntarily onto a hard paper surface. The distributions from B) recorded at 200 mm below the release point corresponded to the same falling distance as those recorded for impact in A). From left to right, top to bottom in each plot: the dotted lines represent model predictions for upright landings (29.3%) and upside down landings (29.3%) at impact, and the expectation for upright vs upside down final landing orientation (50%), if landing orientation were random. (red arrows = dorsoventral axis)

We also compared these data with a probabilistic model that assumed the likelihood of a specimen impacting the surface in a given orientation is proportional to the fraction of solid angles corresponding to how we scored that orientation. Because we used a fixed angle, *θ* = 45 deg, between the horizontal and the body’s cranial-caudal axis to define orientation at impact, this gives a probability of impacting either upright or upside down equal to the solid angle subtended by a spherical cap with polar angle *θ*. This corresponds to a prediction that the fraction impacting the surface upright should be 29.3% = 2*π* (1 – cos *θ*)/4*π* = (1 - cos 45 deg)/2, the same fraction (29.3%) should impact upside down, and the remaining 41.4% impact at all other possibilities combined. The predictions of this model were not consistent with data for nymphs falling on the hard substrate for live or dead/tucked (χ^2^ test, p < 0.0010 and p < 9 × 10^−14^, respectively). For dead/spread, disagreement with the model could not be ruled out (p = 0.37).

While we were unable to release specimens above leaves reproducibly enough to study their impact and landing distributions *per se*, we did film 49 trials in which 15 different nymphs landed successfully on *A. altissima* leaves (1-6 landings/specimen). (Fig. 7A) Those specimens that landed successfully impacted leaflets upright in 49% of cases and upside down 33% of the time. The remaining 18% impacted on a combination of their caudal (6%), side (6%) or head (2%), or clung on impact to the edge of a leaflet (4%). (Fig. 7B)

**Figure 7.**
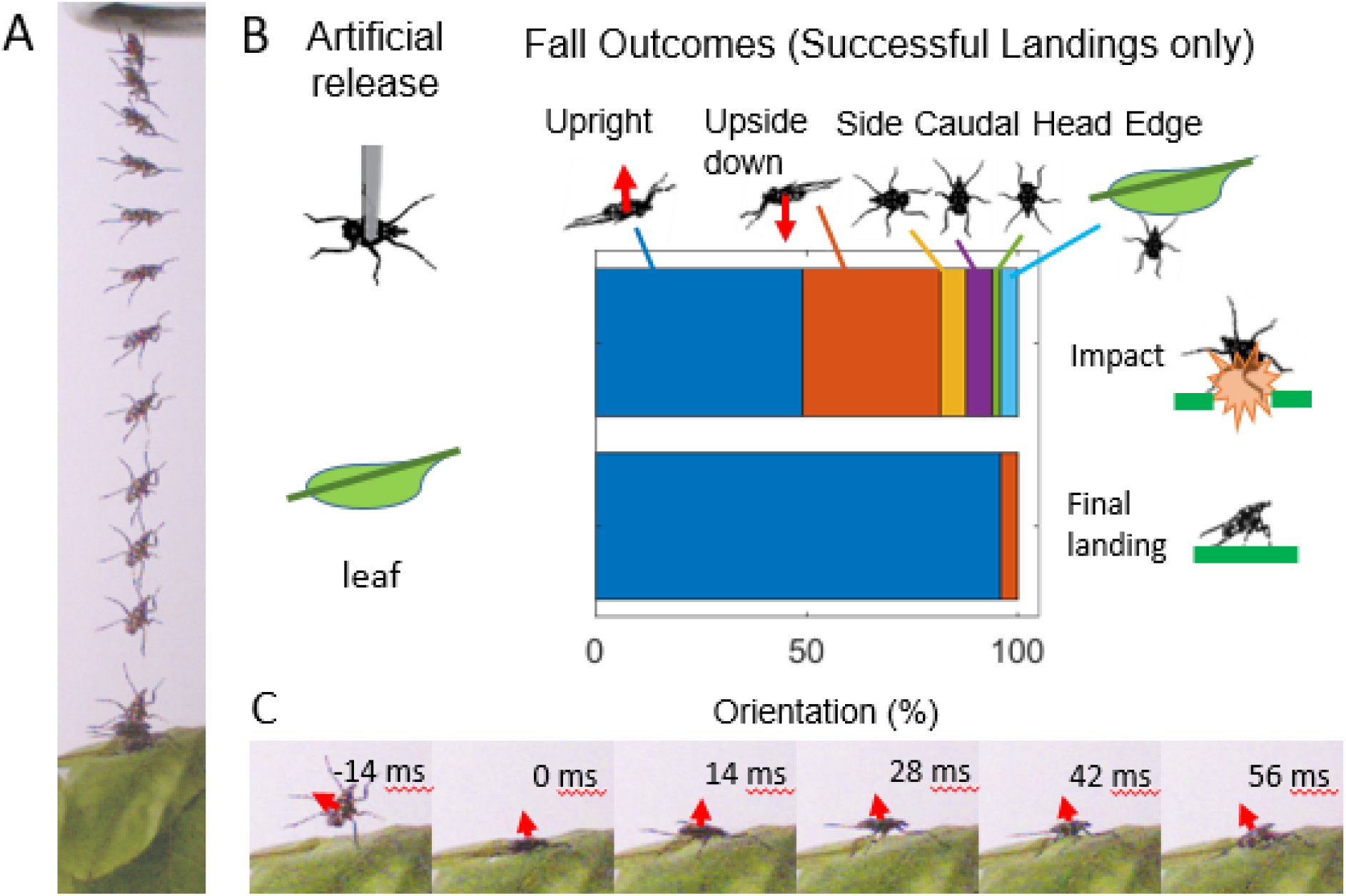
A) Typical image sequence for spotted lanternfly nymphs falling onto *A. altissima* leaflets. B) Orientation distributions at first impact and landing for specimens that successfully landed on leaflets. Because we only characterized successful landings on leaves, these results cannot be compared to the data and models shown in Fig. 6. C) Image sequence showing bouncing from a leaflet. (impact = 0 ms; red arrows = dorsoventral axis)

### 3.4 Bouncing post-impact

Table 2 gives summary statistics for the results of analyzing the post-impact bouncing behaviors. After impact, the vast majority of SLF nymphs bounced at least once, with rebound heights at most a few mm. (Fig. 7C, Fig. 8A) ([S1Movie](https://www.youtube.com/watch?app=desktop&v=Wd-3L8Az7SQ&feature=youtu.be)) As expected from their greater release height, specimens released voluntarily impacted the surface at a significantly higher speed than those released artificially (Tukey’s HSD, p = 0.001); impacts speeds did not differ significantly between any of the other conditions (Fig. 8B). For bounce height, dead/tucked was significantly different from all live conditions (Tukey’s HSD, p < 0.001). The only other significant difference was between voluntarily released live specimens and dead/spread (Tukey’s HSD, p = 0.014). (Fig. 8B, C)

**Table 2.** Summary statistics for spotted lanternfly fourth instar (N4) nymphs impact and post-impact (bouncing) kinematics. (*N* = number of specimens; *n_total_* = total number of trials; *N_bounce_* = number of bounces during landing; *v_imp_* = speed immediately before impact; ***e_c_*** = collisional energy loss). * Only trials in which the nymph landed successfully were analyzed for landing on leaves.

| **Specimen/substrate** | ***N*** | ***n*<sub>total</sub> (n per specimen)** | **Fraction bouncing (%)** | ***N*<sub>bounce</sub>** | ***v*<sub>imp</sub> (m/s)** | **bounce height (mm)** | ***e*<sub>col</sub> (%)** | **Pearson correlation coefficient; linear regression D.F., F statistic, p value** |
| --- | --- | --- | --- | --- | --- | --- | --- | --- |
| Live, voluntary release/ paper | 5 | 15 (2-5) | 100 | 1.07 [0.93, 1.21] | 2.01 [1.91, 2.11] | 3.6 [2.5, 4.6] | 98.3% [97.9, 98.8] | 0.17; 13, 0.38. 0.55 |
| Live/paper | 5 | 31 (6-7) | 93 | 1.02 [0.88, 1.18] | 1.67 [1.62, 1.71] | 4.7 [3.7, 5.6] | 99.0% [98.8, 99.2] | -0.22; 29, 1.5, 0.23 |
| Dead/spread, paper | 5 | 28 (3-7) | 90 | 1.06 [0.85, 1.27] | 1.66 [1.63, 1.69] | 6.6 [5.0, 8.1] | 95.5% [94.5, 96.5] | 0.023; 26, 0.014, 0.91 |
| Dead/tucked, paper | 5 | 39 (5-9) | 93 | 0.78 [0.64, 0.91] | 1.72 [1.68, 1.76] | 8.5 [7.5, 9.5] | 94.4% [93.8, 95.1] | 0.42; 37, 8.1, 0.007 |
| Live/ leaflet* | 8 | 20* (1-5) | 71 | 1.05 [0.89, 1.21] | 1.76 [1.71, 1.82] | 4.2 [3.2, 5.2] | 97.3% [96.8, 97.9] | 0.36; 18, 2.6, 0.12 |

**Figure 8.**
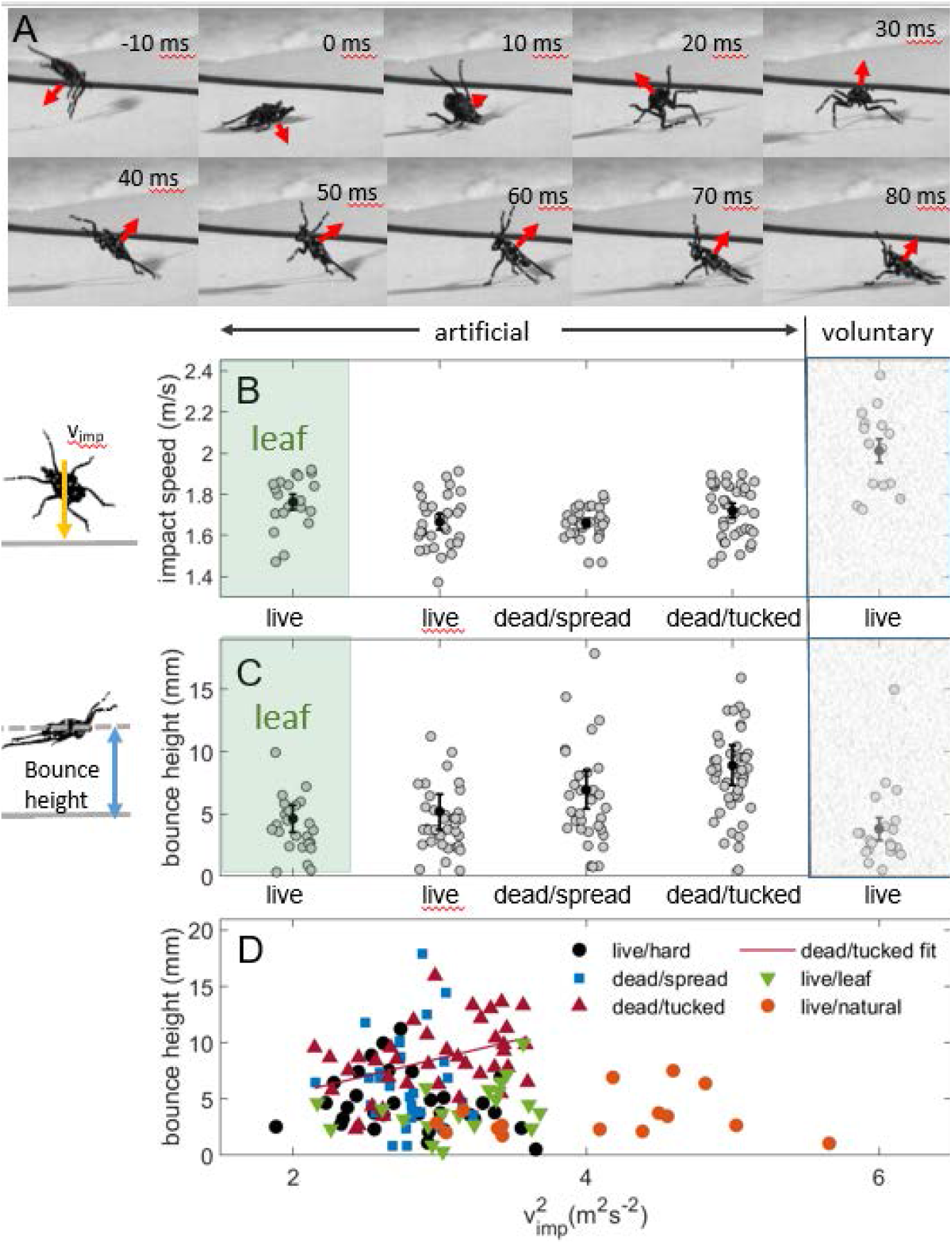
A) Image sequence from a video of fourth instar spotted lanternfly nymph landing on its back on the hard substrate, bouncing, and finally landing upright. (impact = 0 ms; red arrows = dorsoventral axis) Distributions for B) bounce height and C) impact speed, v_imp_, for live and dead specimens artificially dropped from 20 cm onto the hard substrate and leaves and live specimens voluntarily releasing from 35 cm. (black circles: mean; error bars: 95% CI; g circles: all data, jittered for visibility.) D) Bounce height vs 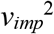 (∝ kinetic energy before impact).

For the 20 trials for which the bounce trajectory could be measured for landing on leaflets, the impact speed and maximum bounce height were consistent with that for live nymphs impacting the hard substrate. (Fig. 8B, C) One notable difference was that the compliant leaflets always recoiled and vibrated after impact. ([S1Movie](https://www.youtube.com/watch?app=desktop&v=Wd-3L8Az7SQ&feature=youtu.be)) For both substrates and all specimen preparations, these bounce heights corresponded to a near-total loss of initial kinetic energy upon colliding with the substrate. Bounce height was weakly correlated with kinetic energy (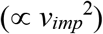) among live specimens, and only dead/tucked specimens had bounce heights that varied linearly with kinetic energy (Fig. 8D). We suspect that the difference in bounce height between live and dead specimens (Fig. 8C) was due in part to the tendency of live, but not dead, nymphs to adhere to the substrate on or immediately after impact with one or more feet. We also observed some high bounces among dead/spread specimens when their legs remained extended on impact, appearing to act as springs rather than collapsing as observed for live specimens.

### 3.5 Final landing behavior

First we consider results for the orientation at final landing (e.g., when the specimen came to rest on the substrate.) A comparison of final landing distributions found that artificially released SLF nymphs were significantly more likely to land upright on the hard substrate when live (56%) than dead with legs spread (35%) or tucked (33%) (Fisher exact test, p = 0.046 and 0.011 respectively). (Fig. 6A) While the number of upright final landing distribution for live specimens released artificially (χ^2^ test, p = 0.180) were consistent with random chance, this was not true for those releasing voluntarily, 100% of which finally landed upright (p = 0.00011). In addition, compared to artificially released specimens, significantly more SLF nymphs releasing voluntarily were oriented upright on final landing (p = 0.00043). (Fig 6B) The fraction of dead specimens that landed upright was lower than predicted by random chance for legs tucked (χ^2^ test, p = 0.0027); for legs spread, the smaller number of observations led to an insignificant test (p = 0.106).

Next, we consider how the orientation at final landing relates to that at first impact. For live nymphs released artificially, the distributions of orientations at impact were significantly different from those upon final landing on the hard substrate (Fisher’s exact test, p < 1.3 × 10^−10^), with a higher percentage achieving an upright orientation at final landing (56%) than on impact (17%). The impact and final landing orientation distributions for voluntary releases also differed significantly (Fisher’s exact test, p = 0.042). The reason why the impact and final landing distributions differed was due in part to the fact that most specimens bounced at least once upon impact and frequently changed orientations as a consequence. Those nymphs that did not land fully upright immediately after bouncing often were able to pull themselves upright as part of a continuous sequence of motion. ([S1Movie](https://www.youtube.com/watch?app=desktop&v=Wd-3L8Az7SQ&feature=youtu.be)) The minority of nymphs that did not bounce on impact either adhered immediately to the substrate upon impact, rolled, or slid to a stop. We observed during preliminary trials that many nymphs that did not come to rest upright upon landing eventually were able to self-right terrestrially without assistance, although we did not study this behavior further.

For landing on leaves, the distribution of orientations also differed significantly between impact and final landing for successful landings (Fisher’s exact test, p < 1 × 10^−7^). The vast majority (96%) of successful final landings were upright, with the remainder oriented upside down. Because we only characterized successful landings on leaves, we did not compare these results to data for all fall outcomes for the other conditions, or to the model, which requires an analysis of all outcomes. SLF nymphs relied on behaviors similar to those found for the hard substrate to cling to leaves after impact. Due to a combination of bouncing, sliding or leaflet vibration, in 33% of the successful landings, the nymphs landed on a surface different from the one on which they made initial impact (i.e., a different leaflet or nearby stem.) In several cases, we observed nymphs grasping a leaflet by its edge by one or more feet and pulling itself successfully onto the surface after a struggle. ([S1Movie](https://www.youtube.com/watch?app=desktop&v=Wd-3L8Az7SQ&feature=youtu.be))

## 4. Discussion

In summary, our study supports the following conclusions: first, spotted lanternfly nymphs falling through ecologically relevant distances used a combination of all righting mechanisms available to them (8), including aerial re-orientation, re-orientation during bouncing, and terrestrial righting, the last of which we do not discuss here. This diverse, flexible arsenal of landing tactics provides SLF nymphs with a variety of ways to respond to surfaces with unpredictable positions, orientations, compliances, textures and other mechanical properties.

Second, our measurements also provide support for SLF nymphs employing both passive and active righting. In virtually all trials, live SLF nymphs assumed a stereotypical falling posture similar to those reported previously for falling pea aphids (11), stick insect instars (2), and geckos (31), as well as gliding ants (9) and spiders (10). On average, they assume this posture within 0.079 s [0.029, 0.129] s after release. Supporting the hypothesis that this posture increases drag, we found that the terminal speed for live and dead specimens with legs spread was significantly lower than for dead ones with legs tucked compactly against the body. Dead specimens with legs tucked also were significantly less likely to impact upright than either live or dead specimens in the falling posture. This supports the argument that the stereotypical falling posture contributes to aerial righting (9,11). When live specimens were able to release voluntarily from surfaces, they were predominantly oriented upright after falling 200 mm to a greater extent than live specimens released artificially. This suggests that when allowed to release voluntarily, SLFs may be modulating initial release conditions to achieve greater upright landing success. The rotational kinematic data during falling indicated that compared to dead specimens, live nymphs rotated more quickly upon release and decreased their rotation rates during falls, likely due to their observed ability to push off the surface of last contact and actively move their legs midair, whereas dead specimens tended to increased their mean rotation rates during falling. Because a nonzero change in rotation rate during the aerial phase requires either a net aerodynamic torque or a change in rotational inertia, these findings suggest likely roles for a combination of aerodynamic torque and active control in determining fall outcomes. Taken together, our kinematic results point to a role for active control to achieve righting during the aerial phase.

Third, our findings indicate SLF nymphs make use of novel active righting behaviors immediately after impact. These motions are distinct from terrestrial righting as previously studied because the nymphs in question enact them before coming to rest on their backs. This interpretation is supported by our finding that, in spite of impacting the surface with similar speeds and orientations, live SLF nymphs finally landed upright significantly more often than dead specimens, even those with legs spread. To understand this phenomenon, we first note that almost all live and dead specimens bounced after impact so as to dissipate most of their pre-impact energy (> 97% for live and >94% for dead specimens), similar to values reported for crash-landing locusts (76%) (24) and cockroaches running into walls (95%) (32). This is important because nymphs benefit from dissipating most of their kinetic energy quickly in order to land securely, while retaining enough kinetic energy to surmount potential energy barriers that can prevent the reorientations required for righting. (7) Consistent with this picture, live SLF nymphs were observed to reorient while rebounding from the substrate, often using grasping or adhesion to the substrate and complex leg motions to lever into a final upright posture.

Collisional energy losses, bounce heights, and subsequent reorientation motions during rebounds were similar for SLF nymphs landing on compliant leaves and hard surfaces. However, several new landing behaviors also were observed, including clinging to the very edge of leaflets, grasping stems and bouncing onto and landing on lower-lying leaflets after initial impact. This was true in spite of the fact that specimens impacted leaves oriented at a variety of angles to the horizontal, and that the leaves recoiled and oscillated on impact. The SLF nymphs’ effectiveness at clinging with a single foot or claw on leaves and their ability to use their arolia for adhesion enhance their ability to settle into a final upright orientation successfully following impact at a variety of angles. This is important because leaflets and other potential perches are encountered at a wide variety of angles in natural habitats.

Taken as a whole, these findings indicate that SLF nymphs falling into underlying foliage could slow down gradually via successive collisions, each of which affords the SLF an opportunity for landing securely. This interpretation is consistent with an earlier study in which pea aphids were induced to drop when on different host plants. The authors found that the probability of dropping pea aphids landing within a host plant instead of on the ground increases approximately linearly with increasing release height (5)--as would be expected if landing success depends on multiple attempts--as opposed to reaching a plateau--as would be expected if the limiting factor was the time required to self-right aerially. Thus, the ability of SLF nymphs to cling securely to the complex foliage of their preferred host suggest that landing upright in itself might not be a necessary or preferred strategy. This possibility deserves to be considered in studying of aerial righting and related phenomena.

Finally, we found that the outcomes of landings after SLF nymphs launched voluntarily from walls were very different from when they were released artificially with minimal speed and rotation rate. Similarly, falling pea aphids were reported to self-right only when released with nonzero initial spin (11). This finding suggests that there is some aspect to preparation or voluntary release that can potentially alter the initial conditions of the fall, setting them up for more upright outcomes. It remains to be explored how the more detailed aerial motions of this species relate to postural control with the goal of ensuring an upright landing. For example, ants (9), spiders (10) and stick insect instars (12) have been shown to use coordinated motions of their legs and appendages during falling to initiate, reorient and stabilize their body orientation.

Taken together, these results point to the importance of studying both aerial and post-impact righting behavior. While most studies on righting behavior during falling have focused on aerial righting, for spotted lanternfly nymphs, post-impact reorientation plays a central role in achieving a final, upright posture in an exceedingly short period of time. Reorientations after impact due to bouncing in particular are so rapid that they cannot be detected without high-speed imaging. The significantly different outcomes observed when specimens were allowed to launch voluntarily from surfaces, combined with some of the unique behaviors observed during falling on leaves, points towards the need to conduct tests in naturalistic environments whenever possible, to better understand ecologically significant behaviors.

## Acknowledgements

We wish to express our appreciation to Eric Beery for work on preliminary planning and pilot experimental studies, Charles Bone and Jennie Ciborowski for help locating specimens and identifying plants in the Haverford College Arboretum, Hongyou Lin for journal article translations, and Naia Hsieh, Sydney Hsieh and Karen Masters for assistance in collecting specimens.

## Data Availability

All data and software required to reproduce the figures and results are included in the Supporting Information.

