## Supplemental Information for "Many ways to stick the landing: novel righting strategies allow spotted lanternfly nymphs to land on diverse substrates"

Spotted lanternfly nymphs use multiple righting behaviors during landing

May 3, 2021

### Supporting Information

**S1 Dataset.** Data and code supplementary information. R-code, Matlab scripts and datasets used to generate the results discussed in the main text.

**Click here to download Dataset 1**

**S1 Movie:** Sample high-speed videos of spotted lanternfly fourth instar nymphs falling, bouncing and landing on a hard paper substrate and leaves.

**Click here to view S1 Movie**

**S1 Table.** Body length and mass of fourth instar spotted lanternfly nymphs. See Fig. 1B for how the body length was defined.

| Body length (mm) | Mass (mg) |
|---:|---:|
| 12.4 | 86.6 |
| 11.7 | 76.5 |
| 12.3 | 66.8 |
| 11.6 | 46.6 |
| 12.1 | 76.0 |
| 11.5 | 51.4 |
| 12.0 | 52.6 |
| 12.4 | 58.4 |
| 11.6 | 76.0 |
| 10.3 | 39.5 |
| 11.0 | 54.6 |
| 12.1 | 68.9 |
| 12.2 | 93.5 |
| 11.9 | 69.8 |
| 12.6 | 100.4 |
| 11.0 | 42.0 |
